# Rewiring vascular patterning through translational control in *Arabidopsis*

**DOI:** 10.64898/2026.01.29.702522

**Authors:** Donghwi Ko, Raili Ruonala, Huili Liu, Ondrej Novak, Karin Ljung, Nuria De Diego, Robert Malinowski, Ykä Helariutta

## Abstract

Plant vascular systems exhibit a wide developmental spectrum, from rigid woody tissues to soft, fleshy storage tissues. We show that increasing polyamine thermospermine transport into wild-type rootstocks, together with cytokinin, reprograms xylem identity from woody to fleshy in *Arabidopsis*. Our findings establish thermospermine as a mobile developmental signal and suggest a strategy for engineering plant vascular architecture.

## Main

Secondary vascular tissue in stems, hypocotyls, and roots mainly comprises phloem, cambium, and xylem, with the cambium functioning as a bifacial stem cell population that produces both phloem and xylem lineages. The phloem contains sieve elements, companion cells, and parenchyma, whereas the xylem consists of vessel elements and parenchyma cells. Across plant species and organs, vascular architecture exhibits remarkable diversity^1–3^. Tree trunks are dominated by vessel-rich xylem but contain relatively few parenchyma cells^1^. In contrast, storage organs such as turnips and radishes develop a fleshy xylem enriched in parenchyma cells, enabling them to store water and resources^2^. These contrasting forms represent distinct developmental trajectories arising from the cambium.

*Arabidopsis thaliana* research has demonstrated that the vascular development is tightly controlled by the interplay of signalling molecules and transcription factors^4^. Auxin signalling induces transcription of the basic helix-loop-helix (bHLH) transcription factors TARGET OF MONOPTEROS 5 (TMO5) and LONESOME HIGHWAY (LHW), which heterodimerise to promote vessel differentiation and induce expression of *LONELY GUY* (*LOG*), which activates cytokinins, and *ACAULIS5* (*ACL5*), a polyamine thermospermine synthase^5,6^. A second bHLH group, SUPPRESSOR OF ACAULIS 51 (SAC51) and its paralogue SAC51-LIKE 3 (SACL3),antagonises TMO5-LHW heterodimerisation and thereby suppresses vessel formation and transcription of *LOG*s and *ACL5* ^5,6^. Translation of *SAC51* and *SACL3* (*SACL*) is repressed by upstream open reading frames (uORFs) in their transcripts, but this repression is relieved by thermospermine^7,8^. Thermospermine thus enhances *SACL* translation, inhibits vessel differentiation and maintains parenchyma identity ^9^. Recent work has uncovered an additional translational regulation layer involving OVERACHIEVER (OVAC), an rRNA methyltransferase that catalyses the m^3^U2952 modification in the peptidyl transferase centre of 25*S* ribosomal RNA. The *ovac* mutant, which lacks m^3^U2952, exhibits reduced translation efficiency for *SACL* and increased translation for *LHW*, resulting in enhanced vessel differentiation and upregulation of *ACL5* transcription and thermospermine biosynthesis^10^.

We performed reciprocal grafting between wild type and *ovac*, and control self-grafts (WT/WT, *ovac*/*ovac*). While WT/*ovac* and control grafts showed no significant non-cell autonomous effects in shoots (Extended Data Fig. 1a), wild-type rootstocks grafted with *ovac* scions exhibited substantially enhanced radial growth of the hypocotyl and hypocotyl/root junction, resembling storage organ development in *Brassicaceae* (Fig. 1a and b). A significant expansion phase started at 7 weeks after grafting, but the bell-shaped mini storage organ became noticeable 5 weeks after grafting. (Fig. 1a, Extended Data Fig. 1b - f). We found a significant reduction in the number of xylem vessels per cross-section area in the mini storage organ (mini-tuber) compared to self-grafted wild type before (Extended Data Fig. 1b - f) and during active cell proliferation (Fig. 1b). We next investigated the nature of the underlying mobile signals promoting fleshy tissue development. In line with that the *ovac* mutant exhibits increased *LOG4* transcription (Extended Data Fig. 1g) as well as increased *ACL5* transcription and thermospermine content^10^, metabolite profiling showed elevated levels of thermospermine and cytokinins in the mini-tuber compared to the wild type (Fig. 1c and d). This suggested that the *ovac* scion overproduces these metabolites, which are then transported to the rootstocks as mobile signals. Grafts between *ovac* scions and *acl5* rootstocks produced the mini-tuber with reduced vessel differentiation per total cross-sectional area compared to the wild-type self-graft (Fig. 1e, Extended Data Fig. 1b - f), supporting a model in which transported thermospermine, rather than locally increased thermospermine biosynthesis, drives mini-tuber formation. Consistently, the translation of *SAC51* and *SACL3* was significantly increased while the translation of *LHW* was downregulated in the mini-tuber compared to wild-type self-grafts (*n* = 4 ∼ 7) (Fig. 2a, b and Extended Data Fig. 2a, b). Grafting *ovac* scions onto *sac51 sacl3* rootstocks neither suppressed vessel differentiation nor promoted cell proliferation, underscoring the necessity of *SACL* translational activation by mobile thermospermine for mini-tuber formation (Fig. 1f and Extended Data Fig. 1b - f). Despite the higher accumulation of thermospermine in self-grafted *ovac* compared to mini-tuber (Fig. 1c), mini-tuber development was not induced in these grafts (Fig. 1a and Extended Data Fig. 1b - f). This aligns with the necessity of OVAC-mediated m^3^U2952 and thermospermine for optimal *SACL* translation and inhibition of *LHW* translation^10^. In the *ovac* self-grafts, the increased thermospermine levels fail to enhance *SACL* translation and suppress LHW translation^10^, thereby hindering mini-tuber formation (Fig. 1a). To further dissect the respective contributions of cytokinin and thermospermine to mini-tuber morphogenesis, we next grafted the cytokinin receptor double mutant *ahk2 ahk3* rootstock with an *ovac* scion. We observed significantly reduced cross-sectional area, yet a continued decrease in vessel differentiation per cross-sectional area (Fig. 1g and Extended Data Fig. 1b - f). This indicates that cytokinin perception is crucial for promoting cell proliferation rather than suppressing vessel differentiation during the mini-tuber formation.

**Figure 1.**
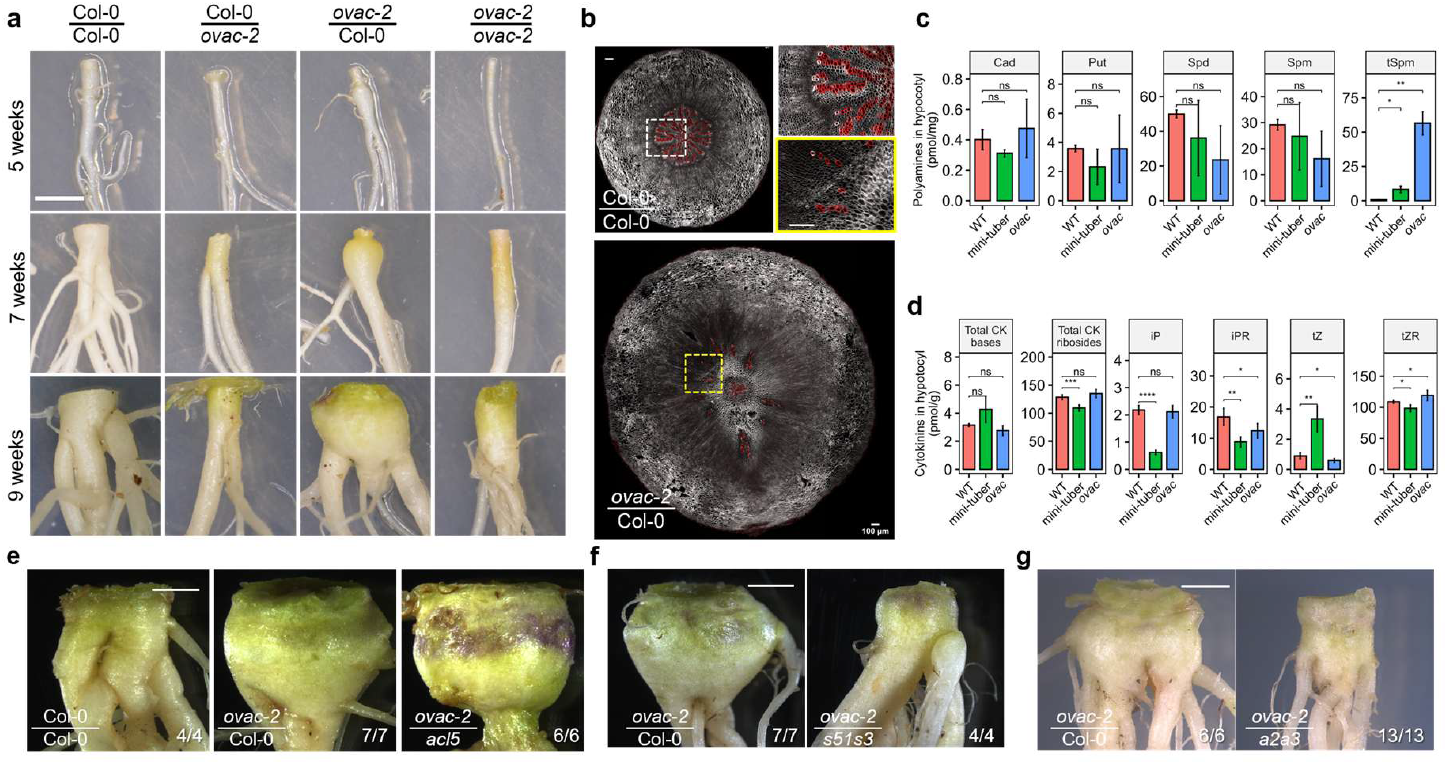
Grafting an *ovac* mutant scion onto wild-type rootstock induces mini-tuber formation. **a**, Photographs of rootstocks of the reciprocal grafts between wild-type and *ovac*-2, and control self-grafts. The grafts were grown under short-day conditions for 5 weeks, 7 weeks, and 9 weeks after grafting (*n* = 2 ∼ 5). Scale bar, 2 mm. **b**, Confocal images of hypocotyl cross-sections of a wild-type self-graft (top panel) and the mini-tuber (bottom panel), taken 8 weeks post-grafting. The cross-sections show SR2200 (grey) and Fuchsin (red) staining. The insets with or without yellow outlines are magnified images of regions marked with white or yellow boxes, respectively, in the hypocotyl cross-sections. Scale bar, 100 µm. **c, d**, Quantification of polyamines (c) or cytokinins (d) in the hypocotyl of *ovac-2* scion and wild-type rootstock grafts (mini-tuber), Col-0 (wild type, WT) and *ovac-2* self-grafts (*ovac*). The grafts for the quantifications were grown under short-day conditions for 10 weeks after grafting (See material and method). Student’s T-test was performed to determine the significant difference (*n* = 3 (c) or *n* = 5 (d). ns, no significance, * *P* ≤ 0.05, ** *P* ≤ 0.01, *** *P* ≤ 0.001, **** *P* ≤ 0.0001). Cad, cadaverine; Put, putrescine; Spd, spermidine; Spm, spermine; tSpm, thermospermine; CK, cytokinins; iP, isopentenyladenine; iPR, isopentenyladenine riboside; *t*Z, *trans*-zeatin; *t*ZR, *trans*-zeatin riboside. **e, f**, and **g**, Images of rootstocks from grafts between *ovac-2* scion with either Col-0, *acl5* (e), *sac51 sacl3* (*s51s3*, f), or *ahk2 ahk3* (*a2a3*, g) rootstocks and Col-0 self-grafts. Grafts were grown under short-day conditions for 12 (e and g) or 11 (f) weeks after grafting. Scale bar, 2 mm.

**Figure 2.**
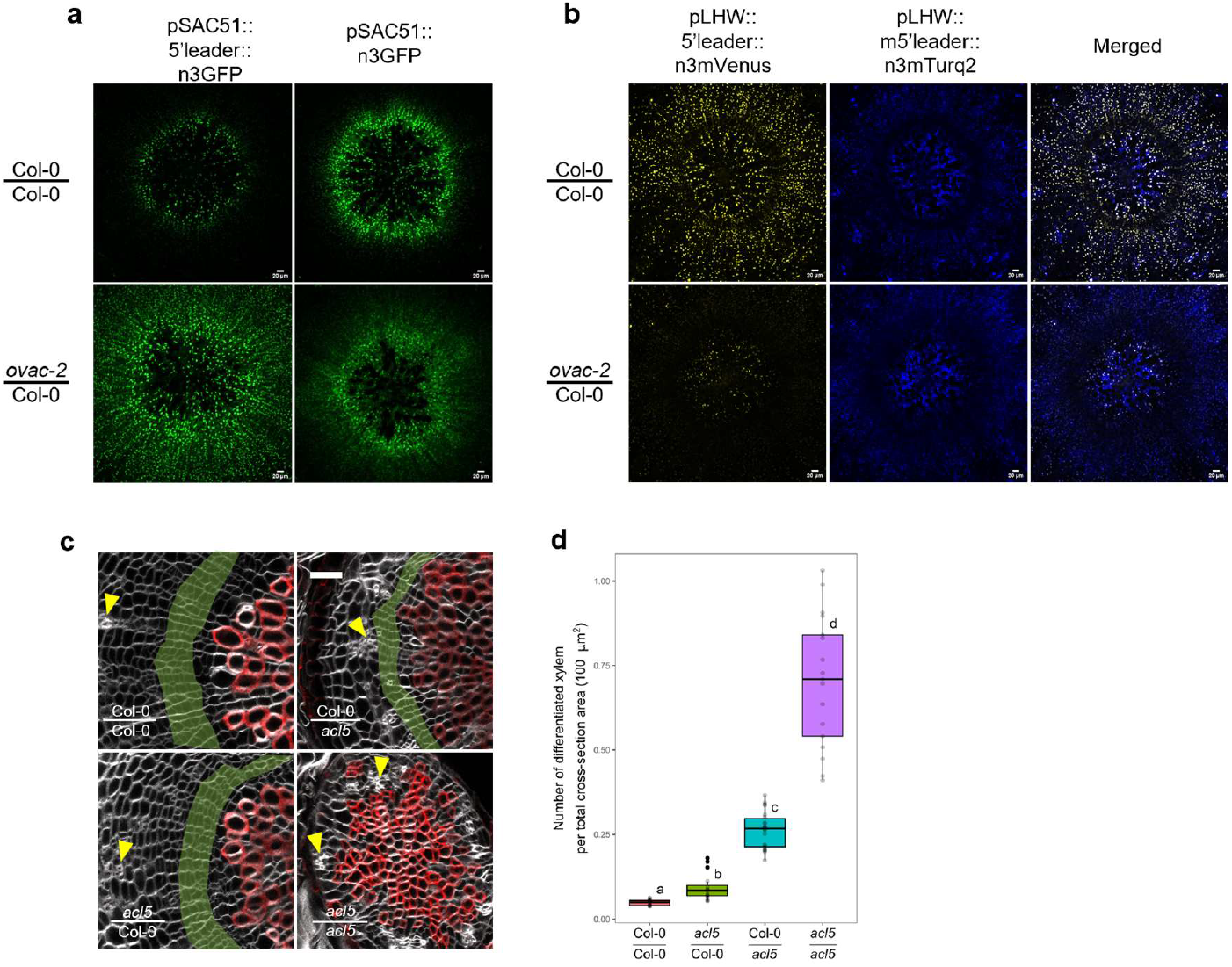
Thermospermine acts as a mobile signal from scions to rootstocks. **a, b**, Translational (*pSAC51::5’ leader::n3GFP* or *pLHW::5’ leader::n3mVenus*) and transcriptional reporters (*pSAC51::n3GFP* or *pLHW::mutated 5’ leader(m5’ leader)::n3mTurquiose2*) for SAC51 (a) and LHW (b) expressed in the rootstocks of 6-week-old grafted plants (*n* = 7 ∼ 9). Each native 5’ leader contains uORFs, whereas in the mutated 5’ leader, the uORF start codons are substituted with stop codons. Scale bars, 100 µm. **c**, Confocal image of hypocotyl cross-sections of reciprocal grafts between wild type and *acl5*, taken 5 weeks post-grafting. Yellow arrowheads indicate phloem sieve elements, and cambial cells are highlighted green. Scale bars, 20 μm. **d**, The number of xylem vessels per cross-section area of grafts shown in **c.** (*n* = 15 ∼ 22, Wilcoxon rank sum test with Bonferroni correction).

Cytokinins are well-known systemic signals^11–13^, but it has not been shown whether thermospermine can also act as a mobile signal in wild-type *Arabidopsis*. To gain further insight into thermospermine transport in wild-type conditions, we performed reciprocal grafting between wild-type and *acl5* plants. Self-grafted *acl5* plants exhibited severely reduced cambial activity, with only a few cells separating the phloem sieve elements and xylem vessels (Fig. 2c, d and Extended Data Fig. 2c - e). This phenotype likely results from premature and excessive vessel differentiation in *acl5*, leading to impaired cambial maintenance and an increased proportion of vessels per total cross-sectional area relative to self-grafted wild-type plants (Fig. 2c, d and Extended Data Fig. 2c - e). In line with previous reports^3^, vessel cell size was also significantly reduced in *acl5* self-grafts (Extended Data Fig. 2f). Grafting wild-type scions onto *acl5* rootstocks partially restored xylem patterning. Specifically, vessel differentiation per total cross-sectional area recovered to 67.2 ± 1.7% of the wild-type level relative to self-grafted *acl5* plants (mean ± s.e.m., *n* = 15 - 22) (Fig. 2c, d; Extended Data Fig. 2c - e). Vessel cell size was also partially restored under these grafting conditions (Extended Data Fig. 2f). In contrast, grafting *acl5* scions onto wild-type rootstocks resulted in a significant increase in vessel differentiation per total cross-sectional area, accompanied by a reduction in vessel cell size compared with wild-type self-grafts (192.8 ± 20.8%; mean ± s.e.m., *n* = 15 - 16) (Fig. 2c, d; Extended Data Fig. 2c–e). Together, these results indicate that thermospermine can be transported from the wild-type scion to the rootstock. However, the amount transported is insufficient to fully rescue the *acl5* phenotype or induce mini-tuber formation. We therefore conclude that thermospermine functions as a non-cell-autonomous signal that promotes *SACL* translation while suppressing *LHW* translation, thereby contributing to cell fate determination and subsequent cytokinin-dependent cell proliferation (Fig. 3).

**Figure 3.**
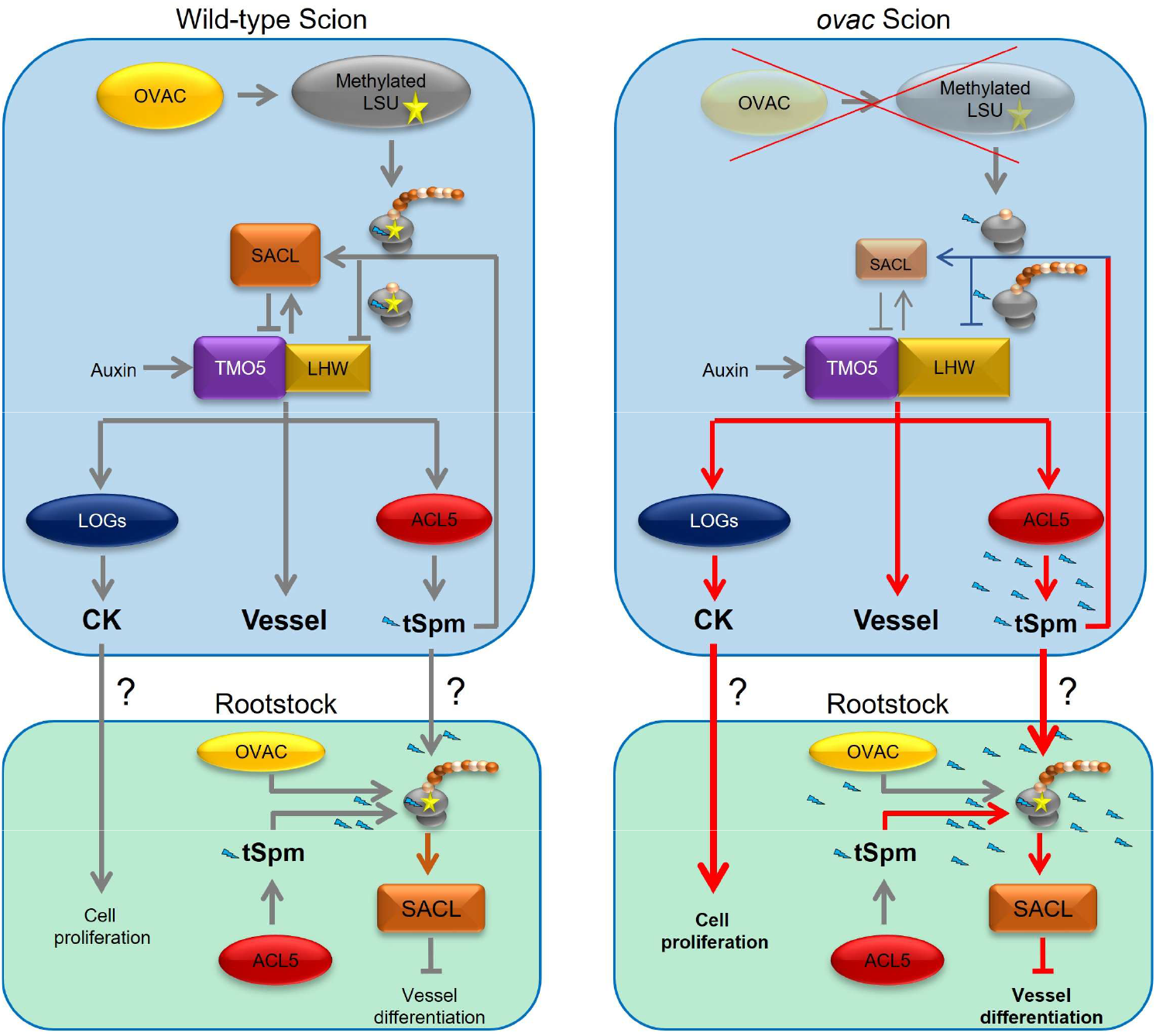
Model for OVAC-thermospermine (tSpm)-mediated control of developmental cell fate decisions in rootstocks. In *ovac* mutant scions (right), the production of cytokinins and thermospermine is increased compared to the wild-type scions, leading to elevated transport of these signals to the rootstock. In wild-type rootstocks, ribosomes carrying the m^3^U2952 rRNA modification, in response to elevated thermospermine transported from the *ovac* scion, promote translation of *SACL* genes while repressing translation of *LHW*, thereby inhibiting vessel differentiation. Concurrently, cytokinins transported from the *ovac* scion stimulate cell proliferation in the rootstock, which, together with thermospermine, drives the formation of a mini-tuber. Red and blue arrows indicate enhanced and suppressed pathways, respectively.

## Discussion

Hormones are signalling molecules that move between cells and bind specific receptors to trigger downstream responses, often at very low concentrations^14^. Thermospermine, which is synthesised by ACL5 in a cell-type-specific manner^15^, is tightly regulated at the level of biosynthesis^16,17^ and accumulates only at low abundance (Fig. 1c). Our grafting experiments demonstrate that thermospermine is transported to rootstocks and acts as a non-cell-autonomous signal that is perceived by m^3^U2952-modified ribosomes (Fig. 1 and 3). Notably, the contrasting vascular architectures generated by different activity levels of the m^3^U2952-thermospermine-SACL-LHW regulatory module closely resemble naturally occurring vascular structures in plants, namely wood and tubers. Wild-type vasculature, characterised by a low parenchyma-to-vessel ratio, is reminiscent of woody tissues, whereas the mini-tuber phenotype, with its high parenchyma-to-vessel ratio, resembles tuberous organs. This parallel suggests that modulation of this translational regulatory module can shift vascular development toward fundamentally distinct organ identities in *Arabidopsis*. This provides an opportunity to investigate whether bona fide storage organs have an enhanced OVAC-thermospermine regulon to favour parenchyma cell identity over vessels.

## Acknowledgements

We thank K. Kainulainen and K. Blajecka for technical support and C. Melnyk for valuable advice on grafting. Y.H. was financially supported by the Center of Excellence in Tree Biology by the Academy of Finland (346139) and Academy Professor by the Academy of Finland (336727). Y.H.’s laboratory was funded by the Gatsby Foundation (GAT3395/PR3); the National Science Foundation Biotechnology and Biological Sciences Research Council grant (BB/N013158/1); University of Helsinki (award 7999920 91), the European Research Council Advanced Investigator Grant SYMDEV (No. 323052). D.K. was funded by an EMBO long-term fellowship ALTF 305-2017. O.N. was supported by the ERC Synergy project ‘Unravelling Spatio-temporal Auxin Intracellular Redistribution for Morphogenesis’ (STARMORPH, reg. no. 101166880). K.L. was supported by grants from the Knut and Alice Wallenberg Foundation (KAW 2016.0352, KAW 2020.0240) and the Swedish Research Council (VR 2021-04938). N.D.D was supported by the project TowArds Next GENeration Crops, reg. no. CZ.02.01.01/00/22_008/0004581 of the ERDF Programme Johannes Amos Comenius. R.M. was supported by the National Science Centre Poland grant (2022/47/B/NZ9/00558).

## Author contributions

Conceptualization: D.K., R.R., Y.H.; Data curation: D.K; Formal analysis: D.K., N.D.D., O.N.; Funding acquisition: Y.H., D.K.; Investigation: H.L., R.R., D.K., F.K.; Methodology: O.N., K.L., N.D.D., D.K., R.R., Y.H.; Project administration: R.R., Y.H.; Resources: K.L., O.N., N.D.D., R.M., Y.H.; Supervision: K.L., N.D.D., R.M., Y.H.; Validation: D.K., R.R., Y.H.; Visualisation: D.K., R.R., Y.H.; Writing – original draft: D.K., R.R., Y.H.; Writing – reviewing & editing: D.K., R.R., Y.H.

## Additional information

The manuscript in press is available from the authors upon reasonable request.

## Materials and Methods

### Plant materials and growth conditions

All *Arabidopsis* mutants used in this study are in the Columbia-0 background (Col-0, wild type, WT). For sterile plant culture, *Arabidopsis* seeds were surface sterilised in bleach (4% sodium hypochlorite with 0.02% tween-20) for 4-5 min with vortexing, or in bleach for 5 min followed by 70% ethanol washes. After the sterilisation, seeds were washed with sterile water for 5-8 times, and stratified in water at 4°C in darkness for 2-4 days. Seeds were plated on half-strength Murashige and Skoog (MS)^18^ medium containing 0.5x MS salts, 1% Difco agar, 1% sucrose and 0.5g/l MES, pH 5.7. The plates were incubated vertically in a 23°C growth chamber with long-day settings (16/8 h light/dark), unless noted otherwise. For experiments requiring an extended period of growth, such as transformation or propagation, the *Arabidopsis* seeds were placed directly on compost (Levington F2) or transferred to compost as seedlings after cultivating on sterile plates for about one week.

### Micrografting

5d or 7d-old seedlings grown under long-day or short-day conditions were grafted following the previously described protocol^19^. Successful grafts were first transferred to half MS media at 7 days after grafting and grown for 7 days before being transferred to soil.

### Polyamine and cytokinin measurements

10-week old grafts of mini-tuber, wild-type and *ovac* self-grafts were used for polyamine and cytokinin quantification. To collect the samples, the shoot part above the graft junction (∼ 0.3 cm from the shoot apex) was removed, and one ∼ 0.5 cm hypocotyl was collected in each tube, and five such replicates were produced of each graft type for cytokinin quantification. 10 ∼ 15 or 3 ∼ 4 hypocotyls for wild-type and *ovac* self-grafts, or the mini-tubers were collected in each tube, and three such replicates were produced for polyamine quantification. For purification and high-sensitivity measurements of the cytokinin species, protocols described in ^20,21^ were followed. Polyamines were purified and measured based on procedures described in^22^,^23^.

### Histology

*Arabidopsis* hypocotyls were fixed in 4% formaldehyde solution in phosphate-buffered saline (PBS) solution for 24 hours and embedded in 4% Agarose for vibratome sectioning following a protocol described in^24^. Transverse sections were made by vibratome Leica VT1200S and stained with 0.1% (v/v) SCRI Renaissance 2200 (SR2200) Stain and 0.05% Fuchsin stain before imaging by confocal microscope. Total cross-section area, xylem area and vessel area were measured using Fiji software^25^.

### Confocal imaging

Confocal images were acquired using a Leica TCS SP8 confocal microscope with objectives 20x/0.70 DRY. The image resolution was set to 512 x 512 or 1024 x 1024 pixels, with a z-step size of 1 µm. Excitation settings were 405 nm (SR2200), 448 nm (mTurquiose2), 488 nm (GFP), 514 nm (mVenus), or 561 nm (Fuchsin staining and mScarlet-I), while emission settings ranged 430 ∼ 445 nm (SR2200), 471 ∼ 495 nm (mTurquiose2), 496 ∼ 521 nm (GFP), 525 ∼ 555 nm (mVenus), and 590 ∼ 615 nm (Fuchsin staining and mScarlet), respectively. To determine xylem vessel numbers per total vascular area, the total vasculature area was measured using Fiji software^25^ and the number of xylem vessels was manually counted based on Fuchsin staining or counted by Cellpose^26^. We measured the cell size of vessels using the Cellpose software^26^ on transverse sections stained with Fuchsin. Within the software, we configured the cell size parameter to 11.5 and employed the Cyto2 model for the vessel cell segmentation.

---

**Extended Data Figure 1.**
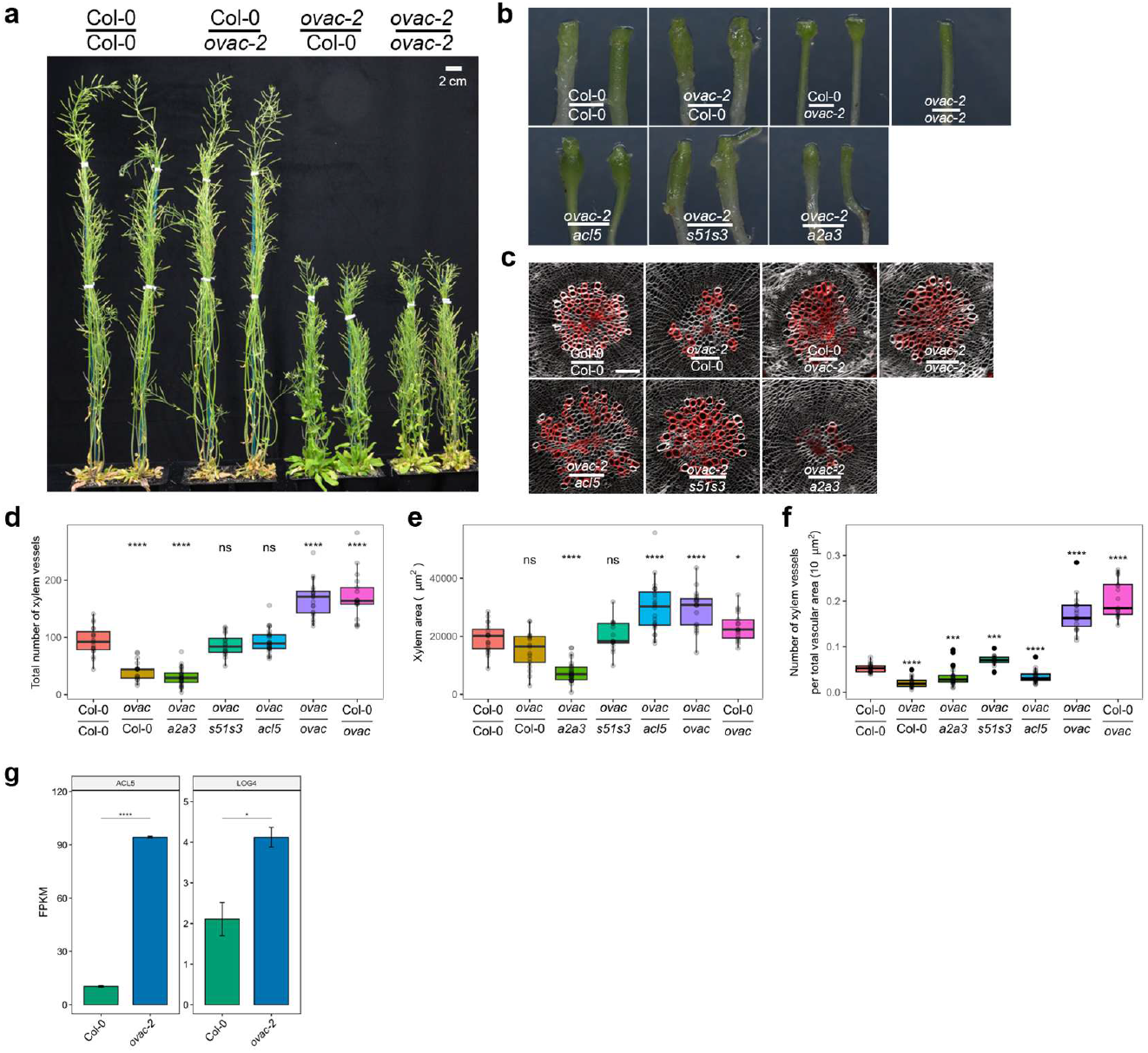
**a**, Photographs of scions of the reciprocal grafts between wild-type and *ovac*-2, and control self-grafts. The grafts were grown under long-day conditions for 8 weeks after grafting (*n* = 15 ∼ 22). Scale bar, 2 cm. **b**, Images of rootstocks from various grafts grown under short-day conditions for 5 weeks after grafting. Scale bar, 1 cm. **c**, Confocal image of cross-sections of hypocotyl from the indicated graft shown in **b**. Grey, SR2200; Red, Fuchsin staining. Scale bars, 50 μm. **d, e** and **f**, The number of xylem vessels (d), total cross-section area (e), or the number of xylem vessels per total cross-section area of grafts shown in c (*n* = 16 ∼ 25, Wilcoxon rank sum test with Bonferroni correction for pairwise comparison). *s51s3, sac51sacl3*; *a2a3, ahk2ahk3*. **g**, Transcript levels of *ACL5* and *LOG4* in wild type and *ovac-2* from transcriptome analysis (*n = 3*). Student’s T-test, ns, no significance, * *P* ≤ 0.05,\*\**P* ≤ 0.01, \*\*\**P* ≤ 0.001. RNA-sequencing data are from ^10^.

**Extended Data Figure 2.**
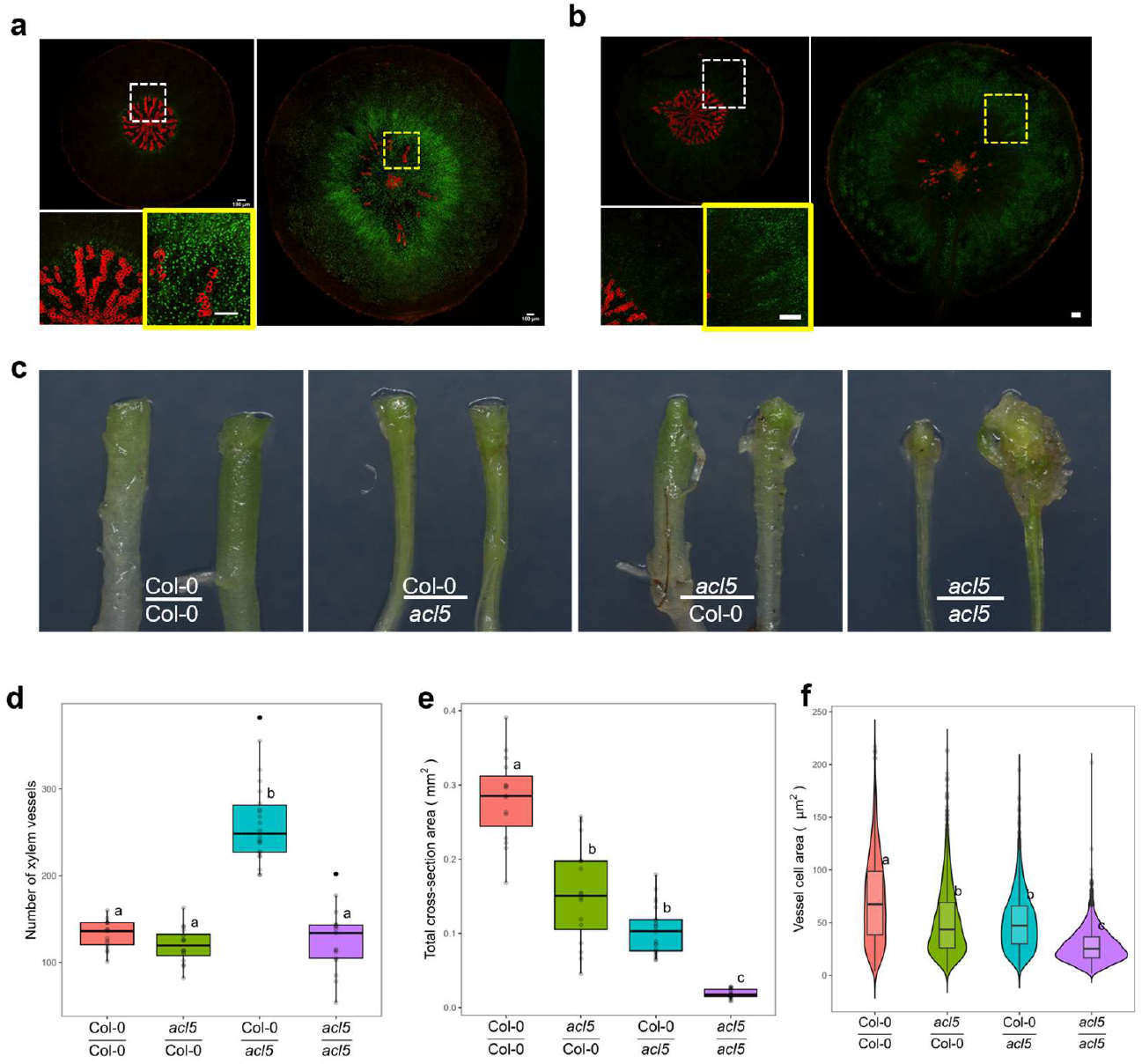
**a, b**, Visualisation of *SAC51* (**a**) and *SACL3* (**b**) translation in wild-type self-graft (left) and the mini-tuber (right). The grafts were grown under the short-day condition for 8 (**a**) or 9 (**b**) weeks after grafting (*n* = 4 ∼ 7). The insets with or without a yellow outline represent magnified images of regions marked with white or yellow boxes, respectively, in the hypocotyl cross-sections. Green, *pSAC51::SAC51 5’ leader::n3GFP* (**a**) or *pSACL3::SACL3 5’ leader::n3GFP* (b); Red, Fuchsin staining. Scale bar, 100 µm. **c**, Images of rootstocks from reciprocal grafts between Col-0 and *acl5* grown under short-day conditions for 5 weeks after grafting. Note that the wild-type self-grafts are the same as those in Extended Data Fig. 1a. Scale bar, 1 cm. **d, e**, The number of xylem vessels (d) or total cross-section area (e) quantified in grafts indicated at 5 weeks after grafting (*n* = 15 ∼ 22, one-way ANOVA and post-hoc Tukey’s test (d) or the Wilcoxon rank sum test with Bonferroni correction (e) for pairwise comparison). **f**, Cell area of vessels in the transverse sections of rootstocks shown in Fig. 2c (*n* = 11 ∼ 17, Wilcoxon rank sum test with Bonferroni correction for pairwise comparison).

